# Predicting microbial community responses to disturbance using genome-resolved trait-based life-history strategies

**DOI:** 10.1101/2025.09.04.674136

**Authors:** Ezequiel Santillan, Soheil A. Neshat, Stefan Wuertz

## Abstract

Understanding how microbial communities respond to disturbance remains a fundamental question in ecology, with broad implications for biodiversity, ecosystem function, and biotechnology. Trait-based approaches offer general rules to predict community responses by linking ecological strategies to measurable traits. While life-history strategy frameworks such as the competitor–ruderal–stress-tolerant (CSR) model are well established in plant and animal ecology, their application to microbial communities has been limited. Here, we experimentally tested how microbial communities shift across a gradient of disturbance frequency in replicated bioreactors treating synthetic wastewater. We applied six conditions by doubling the organic loading rate at different frequencies, from undisturbed to press disturbance, and monitored changes over 42 days using genome-resolved metagenomics, 16S rRNA gene sequencing, biomass quantification, and effluent chemistry. By integrating ordination, network analysis, and machine learning, we identified emergent community-level life-history strategies that aligned with increasing disturbance. These strategies were reflected in functional trade-offs, shifts in community composition, and genomic trait distributions. A simulation-based approach was used to generate a CSR classification of metagenome-assembled genomes, which was consistent with patterns observed in other microbial ecosystems. Our results demonstrate that life-history frameworks can capture predictable dynamics in microbial communities across disturbance regimes. This strategy provides a unifying tool for linking microbial structure, function, and traits across scales, helping to reconcile ecological theory with microbial resource management. More broadly, our findings support the integration of classical ecological theory with microbial genomics to uncover the trait-based principles that govern microbiome function and stability in both natural and engineered ecosystems.

## Introduction

Microbial communities underpin essential functions in both natural and engineered ecosystems, including global biogeochemical cycles [1] and the operation of biotechnological systems [2, 3]. In wastewater treatment reactors, for example, microbial consortia play critical roles in organic matter degradation [4], nutrient removal [5], bioenergy generation [6], and emerging applications such as resource recovery [7] and microbial protein production [8, 9]. However, the stability and performance of these systems depend on how microbial communities respond to environmental disturbances [10], including toxic shocks [11, 12], changes in substrate availability [13–15], temperature fluctuations, or operational disruptions. Predicting such responses remains a major challenge in microbial ecology [16, 17], particularly as many systems function under non-equilibrium and fluctuating conditions [2, 18].

Ecological theory offers powerful tools to understand microbial community responses across environmental gradients [19, 20]. Trait-based approaches, in particular, provide a way to simplify complex dynamics into interpretable axes of ecological strategy [2, 21–23]. Traits, defined as morphologic, physiologic, genomic or phenotypic attributes that affect growth, reproduction, and survival (*i.e*., fitness) [24], can be combined across taxa into community-level indicators known as community-aggregated traits (CATs) [21, 25]. These traits offer mechanistic insights into how microbial communities reorganize their functional potential in response to environmental change [21].

The competitor–stress-tolerant–ruderal (CSR) life-history theory [26] classifies organisms based on how they allocate resources to growth, resource acquisition, or stress tolerance under different combinations of competition, disturbance, and stress. Competitors (C) thrive in stable, resource-rich environments, ruderals (R) rapidly exploit disturbed niches, and stress-tolerants (S) persist under high-stress conditions. Originally conceptualized for plants, the CSR theory has since been adapted to microbes [23, 27, 28] and applied to both laboratory [21] and field studies, especially in soil communities [29–31], using trait-based ordinations and gene-level indicators.

The potential of CSR to explain microbial dynamics in engineered systems was first tested in lab-scale bioreactors [21], which exposed bacterial communities in activated sludge to a gradient of xenobiotic disturbance (3-chloroaniline) and identified life-history strategies based on CATs derived from functional gene profiles. The authors proposed that microbial responses in biotechnological systems could be simplified into ecologically meaningful categories that reflect performance trade-offs. Yet, disturbance is inherently multidimensional [32], and testing other types of disturbance is needed to assess generalizability.

Advances in metagenomics enable trait inference at a higher resolution. While earlier work relied primarily on short-read functional annotations [21], the analysis of metagenome-assembled genomes (MAGs) can provide a more precise characterization of genotypic traits across microbial populations [33, 34]. MAG-based analyses provide a complementary view to amplicon-based structure or short-read function, integrating taxonomy, ecological strategy, and encoded functions. This can enhance the applicability of CSR and other trait-based frameworks in systems where complex interactions and stochasticity co-occur [35, 36]. Further, the recently developed MicroEcoTools package [37] facilitates statistical evaluation of trait-based strategies using metagenomic and functional datasets, promoting reproducibility and accessibility.

In this study, we apply and extend the CSR life-history framework to microbial communities in activated sludge bioreactors subjected to a disturbance gradient created by doubling the organic load at varied frequencies. This disturbance regime influences microbial dynamics by altering competition for oxygen, substrate availability, and spatial niches within the biofilm [36]. Building on prior work [21], we increased experimental replication (n = 5), applied a different disturbance type, and incorporated both 16S rRNA gene amplicon sequencing and genome-resolved metagenomics to track community structure, functional potential, and trait dynamics. Using ordination, network, and machine learning analyses of CATs derived from MAGs and amplicon data, we tested whether distinct life-history strategies emerge along the disturbance gradient. Our goal was to assess the extent to which this classical ecological framework offers predictive insights into microbial trait dynamics and community responses under disturbance, and to support its broader application across both applied and natural microbiome studies.

## Results

### Disturbance shapes bacterial succession, community structure, and functional trade-offs

Bacterial communities differentiated in terms of β-diversity across disturbance levels and with time, as revealed by community analysis through 16S rRNA amplicon sequencing. There was a temporal separation of communities at the undisturbed level 0, the intermediately disturbed levels 1 to 4, and the press-disturbed level 5 as shown by unconstrained ordination (Fig. 1a). This pattern was further supported by constrained ordination at day 42, which emphasized differences across disturbance treatments (Fig. 1b). Multivariate tests yielded significant results for disturbance levels (PERMANOVA-P < 0.001), with no significant effect of heteroscedasticity (PERMDISP-P = 0.86). Variations in β-diversity corresponded with trade-offs in community-level function, as shown by Pearson’s correlation vectors (Fig. 1b), which represent ecosystem function parameters that significantly differed across reactors exposed to different disturbance levels (Table S1). In terms of nitrogen, undisturbed reactors (L0) exhibited the highest effluent nitrate (NO[[-N) concentrations, while press-disturbed reactors (L5) showed the poorest total Kjeldahl nitrogen (TKN) removal. Intermediately disturbed reactors (L1–L4) achieved the most effective TKN removal. Sludge settleability was also highest at intermediate disturbance levels, as indicated by the additive inverse sludge volume index (-SVI). Carbon removal remained consistently high across all reactors (98–99%) and did not differ significantly among disturbance levels. The observed patterns in community diversity and function across disturbance levels resembled those predicted by the CSR framework (Fig. 1c).

**Fig. 1.**
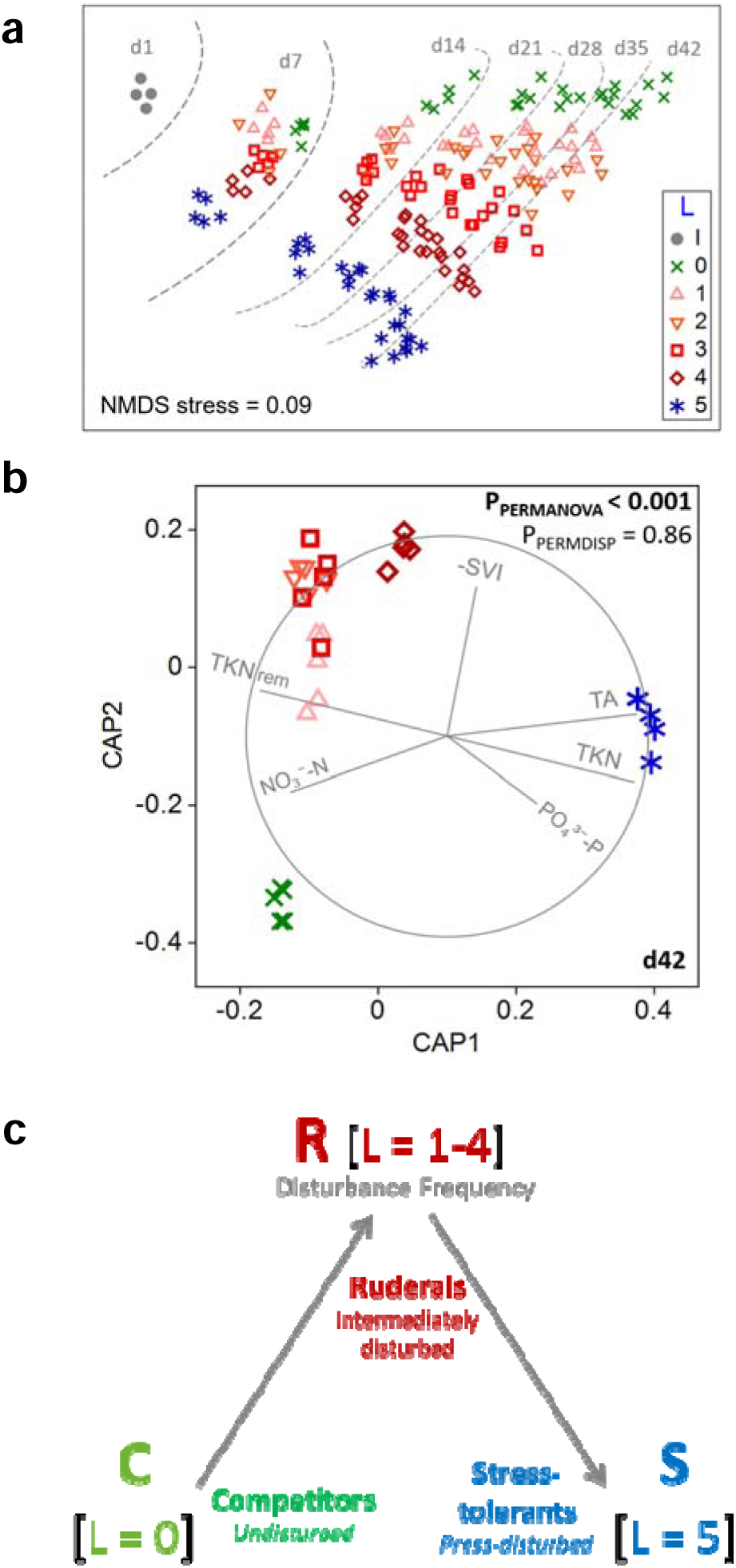
Succession of bacterial communities and functional trade-offs across a disturbance gradient. (**a**) Non-metric multidimensional scaling (NMDS) of 16S rRNA gene amplicon sequence variant (ASV) data across all time points. Bray-Curtis dissimilarities were calculated from square-root transformed abundances. Symbols represent disturbance levels (L0–L5, n = 5 each), and grey dots represent the full-scale plant inoculum (I, n = 4). Dashed lines trace community trajectories from day 1 to day 42. (**b**) Canonical analysis of principal coordinates (CAP) at day 42. Same symbols as panel (a). Vectors indicate Pearson correlations for significantly different (Table S1) effluent and performance parameters: total Kjeldahl nitrogen (TKN), nitrate (NO_3_^-^-N), phosphate (PO_4_^3-^-P), total alkalinity (TA), TKN removal (TKN_rem_), and additive inverse sludge volume index (–SVI). PERMANOVA and PERMDISP p-values indicate significant group separation without dispersion differences. (**c**) Conceptual CSR triangle for d42 bacterial communities showing the dominance of competitor (C), ruderal (R), and stress-tolerant (S) strategies under undisturbed (L0), intermediately disturbed (L1–L4), and press-disturbed (L5) conditions, respectively.

### Genome resolved insights into community clustering and CSR life-history strategies

To gain finer resolution of bacterial dynamics under disturbance, we reconstructed 133 medium- and high-quality MAGs from the day-42 metagenomes (Table S2), following the Minimum Information about Metagenome-Assembled Genomes (MIMAG) framework [38]. Of these, 57 MAGs were classified as having substantial (90–99.9%) or perfect (100%) completeness, and 105 MAGs had low (5–9.9%) or no (0%) contamination, ensuring high confidence in downstream trait and taxonomic analyses. Summary data and CSR classifications for all MAGs are available in Supplementary Table S3. Canonical analysis of principal coordinates (CAP), based on Bray-Curtis dissimilarities of MAG relative abundances derived from coverage, revealed distinct clustering patterns corresponding to specific disturbance levels (Fig. 2a), with significant group separation (PERMANOVA P < 0.001) and no dispersion bias (PERMDISP P = 0.10). A SparCC correlation network of MAGs revealed associations aligned with CSR categories, as indicated by node modularity classes. Although MAGs represent a defined subset of the microbial community, the observed patterns align with and complement the broader 16S rRNA-based ASV data shown in Fig. 1.

**Fig. 2.**
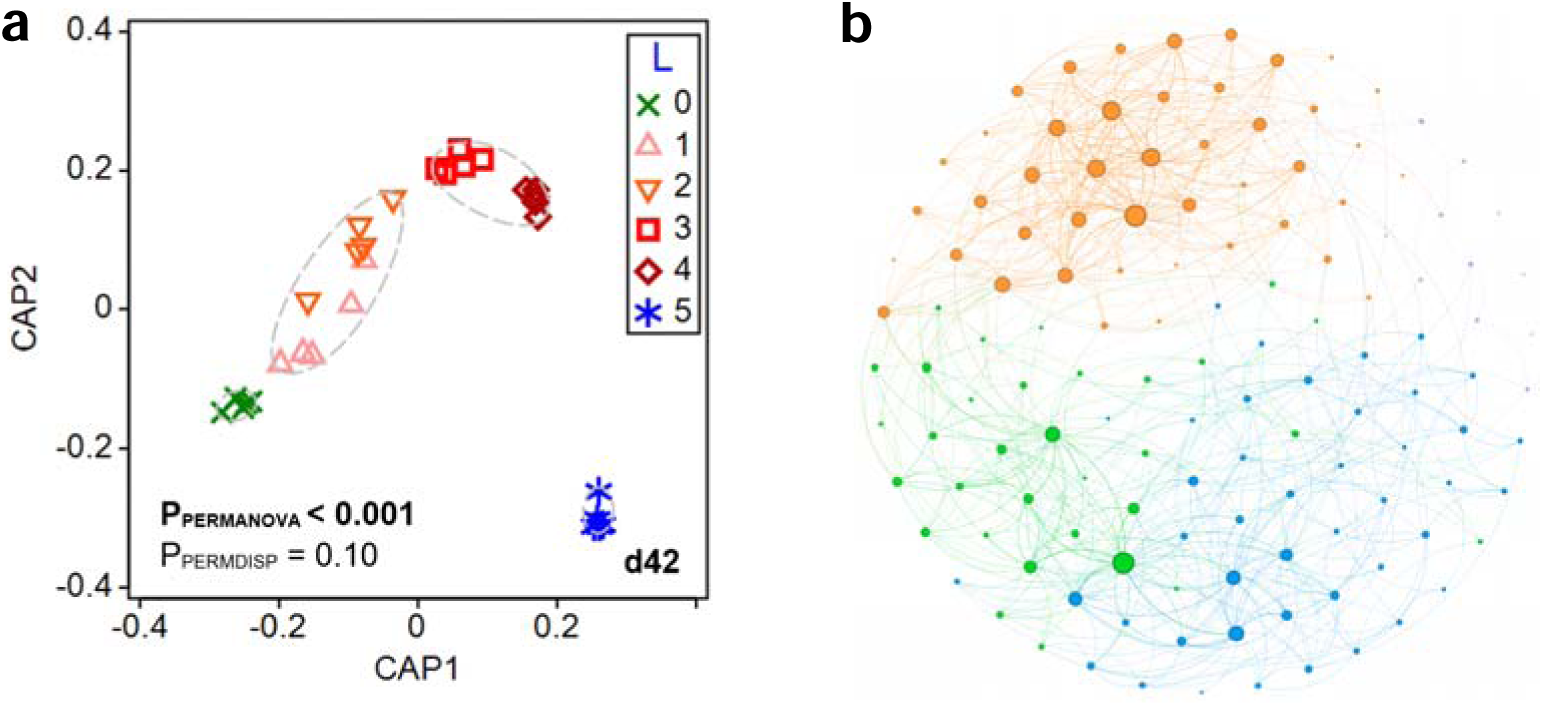
Clustering of bacterial community structure on day 42 based on metagenome-assembled genomes (MAGs). (**a**) Canonical analysis of principal coordinates (CAP) based on Bray-Curtis dissimilarities of square-root transformed relative abundances from 133 medium- and high-quality MAGs. Symbols represent disturbance levels (L0–L5, n = 5). Dashed ellipses represent 85% similarity thresholds from group-average clustering. PERMANOVA and PERMDISP p-values indicate significant group separation without dispersion bias. (**b**) SparCC-based correlation network showing strong positive associations among MAGs (r_adj_ ≥ 0.20). Edge thickness indicates correlation strength, and node size reflects degree. Nodes are coloured by modularity class, corresponding to dominance in undisturbed (L0, green), intermediately disturbed (L1–L4, orange), or press-disturbed (L5, blue) reactors. This analysis represents a MAG-defined subset of the microbial community; broader patterns across all taxa are shown in Fig. 1 using 16S rRNA ASV data.

We further applied the MicroEcoTools *CSR_assign* function to classify the MAGs into CSR life-history strategies (Fig. 3). Of the 133 MAGs, 97 received unique assignments: 18 as competitors (C), 28 as ruderals (R), and 29 as stress-tolerants (S), with several overlapping or unassigned (Fig. 3b). Representative MAGs within each category exhibited consistent relative abundance patterns across the disturbance gradient, with competitor MAGs dominating undisturbed reactors, ruderal MAGs enriched at intermediate disturbance, and stress-tolerant MAGs prevailing under continuous disturbance (Fig. 3a). Genome size distributions generally aligned with CSR expectations: competitors had the largest and least variable genome sizes, ruderals had smaller genomes, and stress-tolerant MAGs spanned a wider range. However, these differences were not statistically significant (Welch’s ANOVA *p* = 0.35; Fig. 3c).

**Fig. 3.**
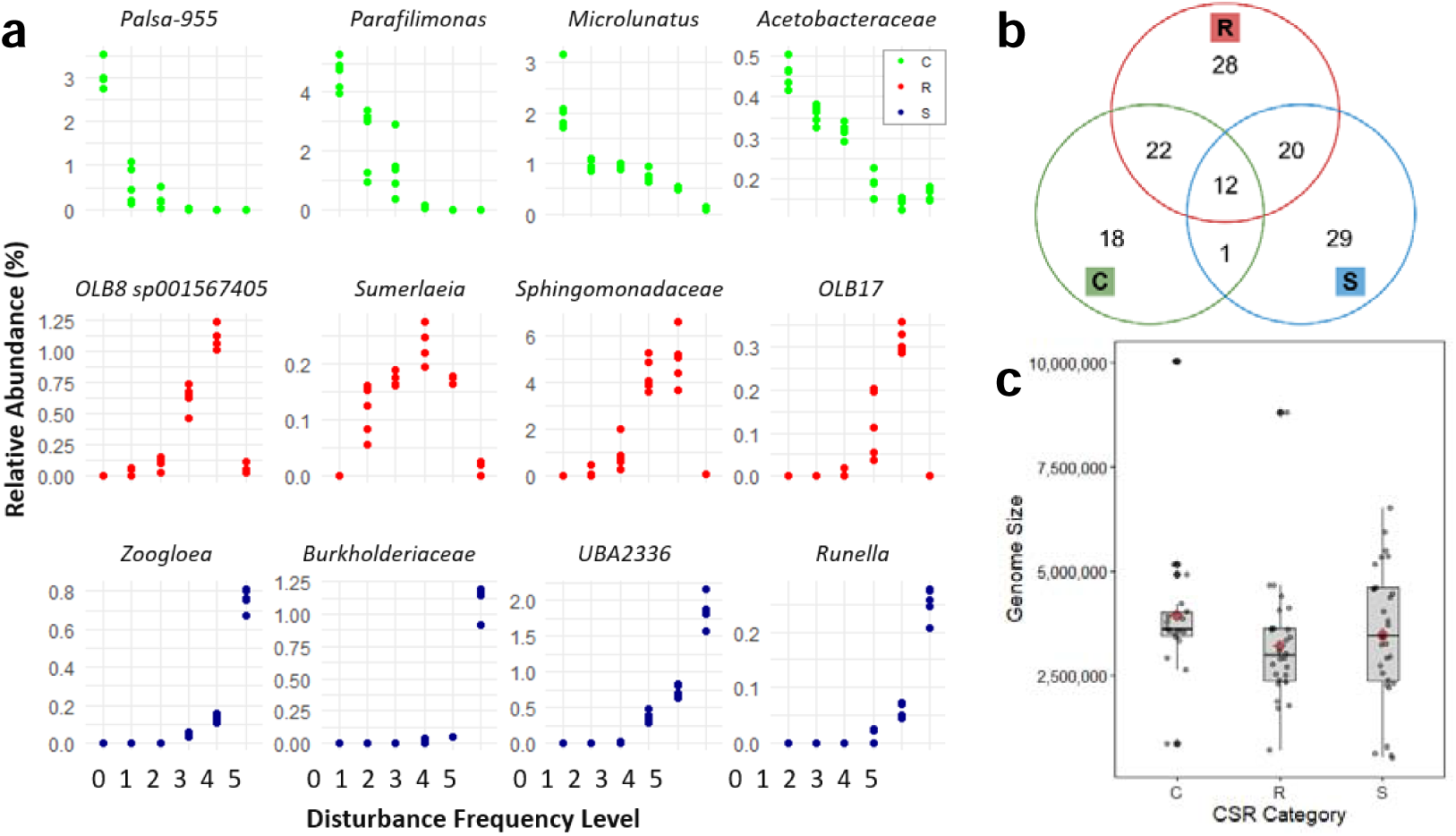
CSR classification of metagenome-assembled genomes (MAGs) and genomic features using the *CSR_assign* function in MicroEcoTools [37]. (**a**) Top four MAGs assigned to each life-history strategy, with competitors (C) in the top row, ruderals (R) in the middle row, and stress-tolerants (S) in the bottom row, based on their relative abundance across disturbance frequency levels. (**b**) Venn diagram of CSR assignments among 133 medium- and high-quality MAGs, indicating overlaps and unique classifications. (**c**) Distribution of estimated genome sizes (in base pairs) for MAGs assigned to CSR categories. Red diamonds display mean values (Welch’s ANOVA p = 0.35). The box bounds the interquartile range (IQR) divided by the median, and Tukey-style whiskers extend to a maximum of 1.5 times the IQR beyond the box.

### Linking microbial genotypic traits to CSR strategies through trait-based analysis

Trait-based analysis of MAGs revealed distinct genotypic signatures associated with different life-history strategies along the disturbance gradient (Fig. 4a). By aggregating Clusters of Orthologous Groups (COG) functional categories across the 133 high- and medium-quality MAGs, we observed clear patterns of enrichment and reduction that aligned with competitor (C), ruderal (R), and stress-tolerant (S) strategies. For example, traits linked to biosynthetic and metabolic efficiency such as secondary structure [Q] and lipid metabolism [I] were enriched under undisturbed (L0) conditions, consistent with a competitor strategy favouring stable environments. In contrast, ruderal-associated traits like transcription [K] and inorganic ion transport [P] peaked at intermediate disturbance levels (L1–L4), where faster metabolic activation and turnover would be advantageous. Stress-tolerant traits, including nucleotide metabolism [F], DNA repair [L], and signal transduction [T], were enriched under press-disturbance (L5), reflecting an adaptation to persist in resource-limited or inhibitory environments.

**Fig. 4.**
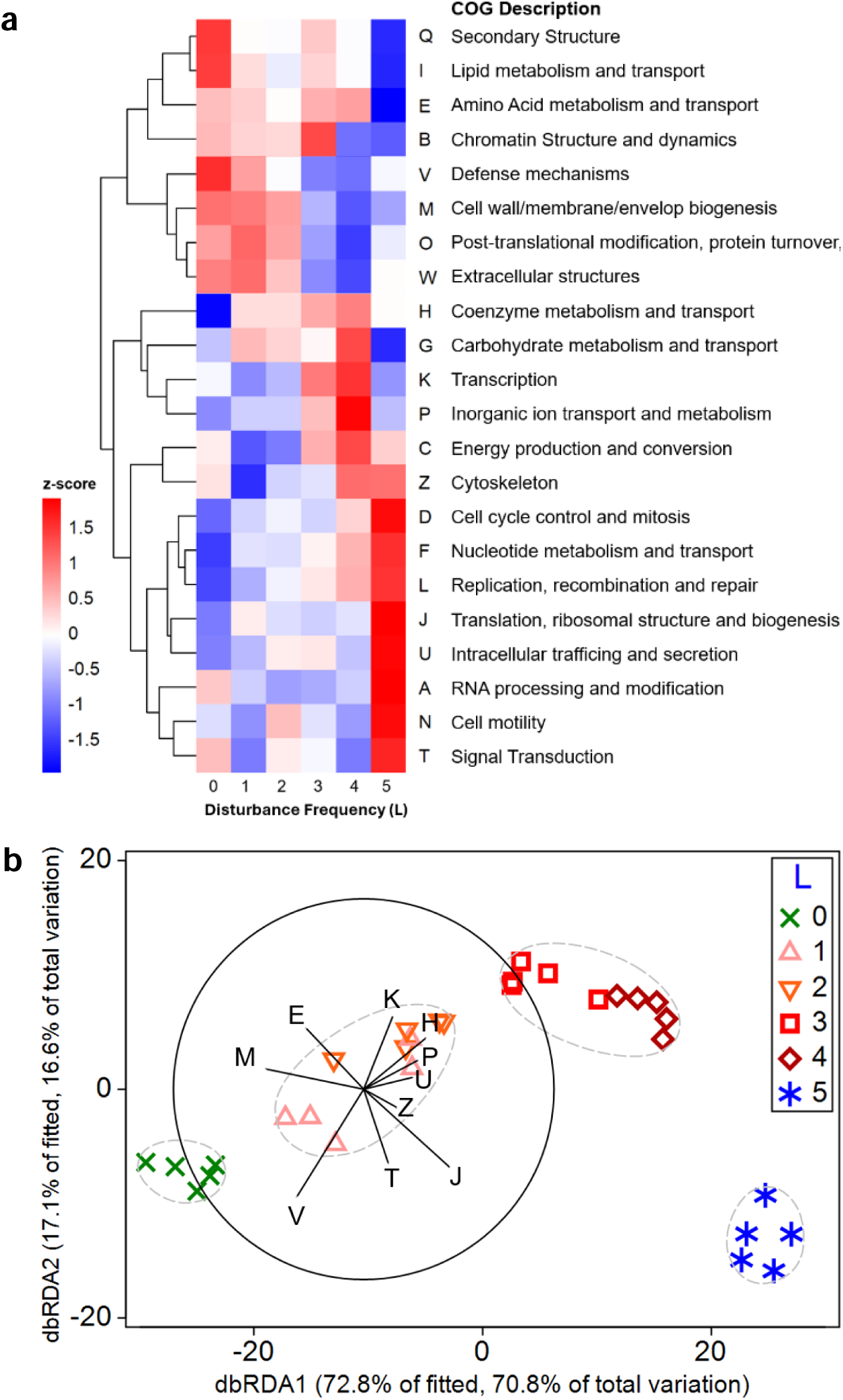
Disturbance frequency shapes bacterial genotypic trait composition across CSR strategies. (**a**) Z-score normalized abundances of Clusters of Orthologous Groups (COG) trait categories across disturbance levels (L0 to L5), derived from 133 medium- and high-quality metagenome-assembled genomes (MAGs) recovered on day 42. COG categories are hierarchically clustered (average linkage). Red and blue indicate relative enrichment and reduction, respectively. Patterns reflect ecological selection of traits associated with competitor (C), ruderal (R), and stress-tolerant (S) strategies. (**b**) Distance-based redundancy analysis (dbRDA) illustrates how variation in COG trait composition explains community structure across disturbance levels. Bray-Curtis dissimilarities were calculated from square-root transformed MAG abundance data. Predictor selection was performed using distance-based linear modeling (DistLM) with stepwise variable selection and AICc minimization, a machine-learning approach that identifies the most explanatory trait complexes. Predictor vectors represent Pearson correlations of model-selected COG categories. Symbols denote disturbance levels (L0 to L5; n = 5 per group), with 85% similarity ellipses based on group-average clustering.

To enable interpretable prediction of how trait composition relates to community structure, we applied distance-based linear modelling (DistLM), a machine-learning approach that performs stepwise model selection and variable ranking based on the corrected Akaike Information Criterion (AICc). From a total of 19 normalized COG trait categories considered, DistLM identified a 10-variable model that best explained variation in the MAG-based Bray-Curtis community dissimilarities (AICc = 109.15), accounting for 97.2% of the fitted variation. The resulting distance-based redundancy analysis (dbRDA) showed that the first two ordination axes explained 70.8% and 16.6% of the total variation, respectively (Fig. 4b). Predictor vectors such as defense mechanisms [V], transcription [K], and translation, ribosomal structure and biogenesis [J] showed strong correlations with ordination axes and revealed consistent clustering of communities by disturbance level, reinforcing the idea that bacterial communities are shaped by trait-linked processes consistent with CSR framework transitions.

Similarly to the MAG-level classification, we applied the MicroEcoTools *CSR_assign* function to assign COG trait categories to CSR strategies based on their enrichment patterns across the disturbance gradient (Table 1). Traits exclusive to each CSR category, such as lipid metabolism [I] for competitors, transcription [K] for ruderals, and DNA repair [L] for stress-tolerants, highlighted distinct ecological trade-offs. Multifunctional traits, including defense mechanisms [V] and cell wall biogenesis [M], appeared across categories (e.g., CS or CR), suggesting context-dependent roles that span resource competition, rapid growth, and stress endurance. These findings support the interpretation that microbial communities can be mapped onto predictable life-history strategies based on their aggregated genotypic potential.

**Table 1.**
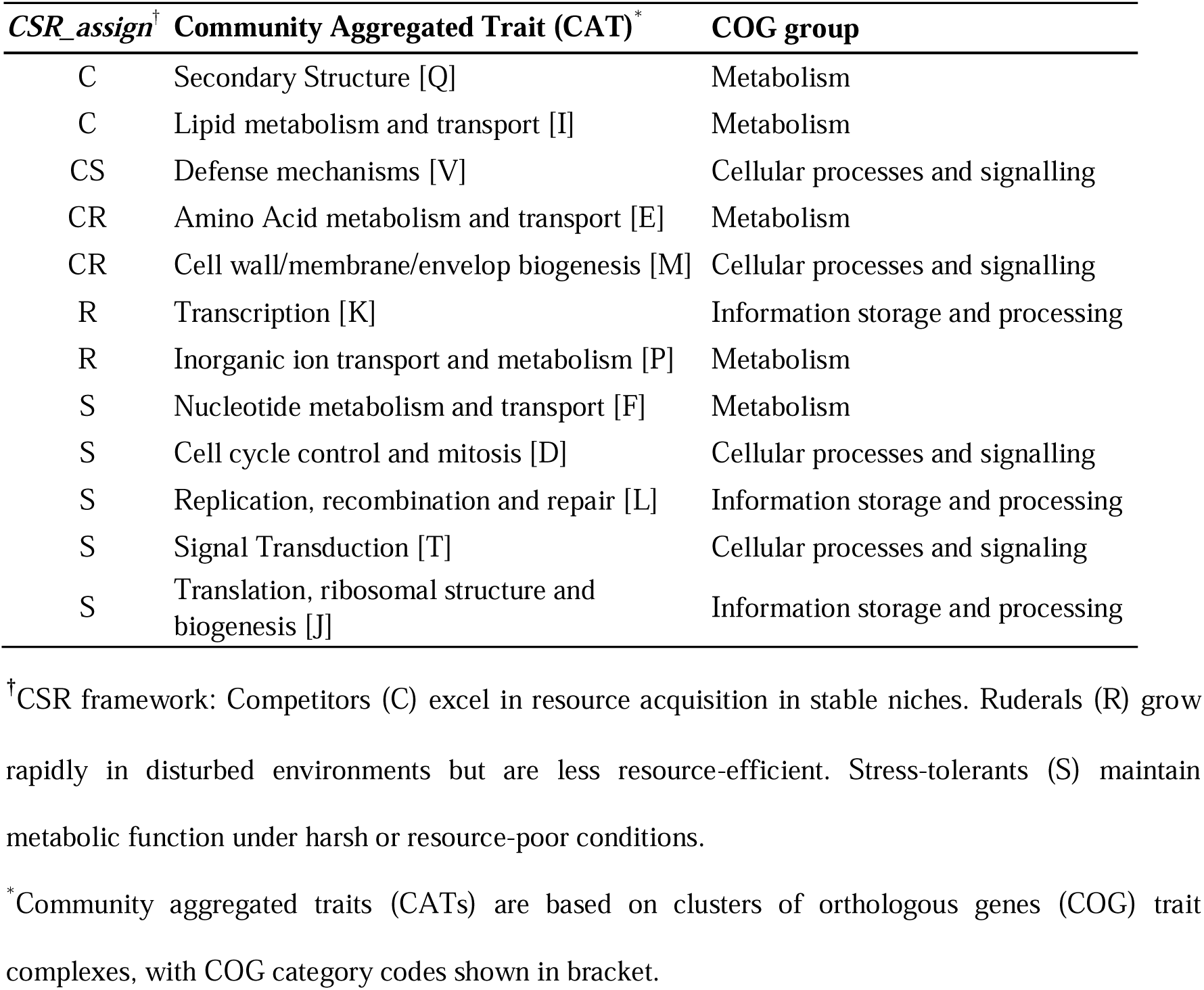
CSR classification of genotypic traits using the *CSR_assign* function in MicroEcoTools [37], based on 133 medium- and high-quality MAGs from day-42 bioreactor metagenomes.

| *CSR_assign*<sup>†</sup> | Community Aggregated Trait (CAT)<sup>*</sup> | COG group |
| --- | --- | --- |
| C | Secondary Structure [Q] | Metabolism |
| C | Lipid metabolism and transport [I] | Metabolism |
| CS | Defense mechanisms [V] | Cellular processes and signalling |
| CR | Amino Acid metabolism and transport [E] | Metabolism |
| CR | Cell wall/membrane/envelop biogenesis [M] | Cellular processes and signalling |
| R | Transcription [K] | Information storage and processing |
| R | Inorganic ion transport and metabolism [P] | Metabolism |
| S | Nucleotide metabolism and transport [F] | Metabolism |
| S | Cell cycle control and mitosis [D] | Cellular processes and signalling |
| S | Replication, recombination and repair [L] | Information storage and processing |
| S | Signal Transduction [T] | Cellular processes and signaling |
| S | Translation, ribosomal structure and biogenesis [J] | Information storage and processing |

<sup>†</sup>CSR framework: Competitors (C) excel in resource acquisition in stable niches. Ruderals (R) grow rapidly in disturbed environments but are less resource-efficient. Stress-tolerants (S) maintain metabolic function under harsh or resource-poor conditions.
<sup>\*</sup>Community aggregated traits (CATs) are based on clusters of orthologous genes (COG) trait complexes, with COG category codes shown in bracket.

## Discussion

This study shows that microbial communities exposed to varied frequencies of organic loading disturbance in bioreactor systems develop distinct configurations of taxonomic composition, functional attributes, and genotypic traits, shaped by both disturbance frequency and temporal dynamics. These patterns align with classical CSR life-history strategies, extending their applicability to microbial ecosystems. These findings build on and validate earlier work [21], which posited that a broad disturbance gradient could represent different ecological conditions within the CSR framework. Specifically, we assumed that undisturbed environments correspond to high competition, where communities are dominated by taxa well adapted to prevailing conditions; intermediate disturbance levels impose strong disturbance pressures that favour ruderals; and continuous (press) disturbance represents high-stress conditions that select for stress-tolerant strategies [39]. By integrating taxonomic profiles, ecosystem function metrics, and CATs derived from both amplicon and genome-resolved metagenomic data, we show that disturbance frequency consistently selects for distinct life-history strategies: competitors under stable conditions, ruderals under intermediate disturbance, and stress-tolerants under press disturbance. These CSR-aligned patterns were evident across beta-diversity, functional trade-offs, trait distributions, and positioning within the CSR triangle.

A key difference from our previous study is the use of metagenome-assembled genomes (MAGs) to derive genotypic CATs, offering finer resolution and stronger linkages between traits and taxa. While MAGs do not capture all functional potential present in the community, they provide organism-resolved insights that complement broader taxonomic or gene-centric approaches. Analyses of genes related to biosynthesis, resource use, and stress response supported CSR assignments, reinforcing the ecological interpretation. These results underscore the value of genome-resolved trait inference for predictive microbial ecology [34] and provide a convergent application of CSR life-history strategies in engineered microbiomes, aligning with similar trait-based interpretations in soil ecosystems [29, 30].

Among traits assigned to a single CSR category, patterns generally matched expected ecological trade-offs. Competitor (C) traits such as secondary structure [Q] and lipid metabolism [I] were enriched in undisturbed L0 reactors, where nitrogen removal was highest. These traits likely support efficient resource acquisition and biosynthesis in stable environments, consistent with prior findings from a study that used 3-chloroaniline (3-CA) as a xenobiotic disturbance in lab-scale activated sludge bioreactors [21]. That study, which applied the CSR framework using CATs from read-based functional gene profiles, also observed biosynthetic trait enrichment under undisturbed conditions. However, L0 reactors in the present study showed poorer settling capacity, suggesting that C traits alone do not guarantee optimal system function. At intermediate disturbance levels (L1 to L4), ruderal (R) traits such as transcription [K] and inorganic ion transport [P] were enriched, coinciding with enhanced TKN removal and settling capacity. These traits likely support fast metabolic responses suited to fluctuating conditions, and their presence aligns with trait combinations observed in the intermediate regimes of both the 3-CA study and recent CSR applications in soils [29], reinforcing the cross-system relevance of this ecological framework. Stress-tolerant (S) traits including nucleotide metabolism [F], cell cycle control [D], DNA repair [L], signal transduction [T], and translation [J] were enriched in L5 reactors, where nitrogen removal was lowest, consistent with survival and maintenance under prolonged stress. This aligns with the broader view that S traits emphasize repair, persistence, and stability under constraint [31, 40].

Distance-based Linear Modelling (DistLM), a machine learning approach, added predictive power and interpretability to our trait-based findings. By identifying a minimal yet optimal set of genotypic traits that best explained variation in MAG-defined community composition, the model validated CSR assignments through independent variable selection. The model-selected traits, such as defense mechanisms [V], transcription [K], and translation [J], contributed strongly to ordination axes and mirrored CSR-aligned clustering patterns. This demonstrates the potential of ML approaches like DistLM to uncover trait–community structure relationships that go beyond correlative descriptions. Rather than merely recognizing patterns, such techniques help disentangle the ecological processes that underlie microbial community dynamics across disturbance gradients. When integrated with hypothesis-driven experimentation and ecological theory, machine learning provides a framework for generating testable predictions and actionable strategies, thereby advancing predictive microbial ecology in both natural and engineered systems [2, 41–43].

Traits assigned to multiple CSR categories showed interpretable but more variable patterns. CR traits like amino acid metabolism [E] and cell wall biogenesis [M] were enriched at intermediate disturbance, suggesting roles in both growth and structural resilience. This is consistent with increased biosynthesis and membrane-related traits reported under moderate disturbance in other systems, including soils [29]. Defense mechanisms [V], classified as CS, were moderately enriched across levels, reflecting functions relevant to both stress buffering and resource competition. Translation [J] appeared in both L0 and L1 to L4 reactors, similar to the 3-CA study where ribosomal traits were shared by competitor and ruderal communities [21]. Some traits diverged from expectations. Lipid metabolism [I], although classified as a C trait, may also contribute to stress tolerance through membrane stabilization, illustrating potential trait multifunctionality. Signal transduction [T], enriched under press disturbance, may mediate rapid environmental sensing, supporting roles in both R and S contexts. Amino acid metabolism [E] also appeared in L0, reinforcing its competitor function noted in prior studies. These mixed patterns align with findings from soil systems where traits do not always map cleanly to CSR categories and can shift functionally with context [29, 44]. This highlights the importance of refining trait classification empirically, particularly in complex and engineered environments.

Although genome size did not differ significantly across CSR categories, the observed trends align with ecological expectations. MAGs classified as competitors had the largest average genome sizes and the narrowest spread, consistent with metabolic versatility and regulatory capacity suited for stable environments where resource efficiency is favored [44]. Ruderal MAGs showed the smallest genomes with constrained variability, which may reflect selection for rapid replication and low biosynthetic burden under fluctuating disturbance. Stress-tolerant MAGs displayed the broadest genome size distribution, potentially capturing a mix of streamlining for persistence and larger genomes carrying diverse stress-response and repair functions. This heterogeneity matches prior findings that different ecological strategies can be associated with both genome expansion and reduction, depending on selective pressures [40]. While genome size alone does not determine strategy, its distribution may reflect underlying trade-offs in growth, regulation, and survival, supporting its complementary role in trait-based CSR classification. However, genome size estimates derived from MAGs have inherent limitations. Even MAGs reported as 100 percent complete by tools such as CheckM may lack substantial genomic regions, since completeness is assessed based on the presence of marker genes [45]. Assuming the bias in genome size estimation is randomly distributed across all MAGs, comparative trends can still offer meaningful ecological insights.

While the focus of this study is on traits, several MAG-assigned taxa align with their CSR classifications based on known biology. Among competitors (C), *Microlunatus* are polyphosphate-accumulating organisms known to store internal phosphorus reserves, supporting stable nutrient removal under undisturbed conditions [46]. Members of *Acetobacteraceae* oxidize low concentrations of simple carbon compounds and can contribute to stable biofilm communities, favouring resource-efficient niches. *Parafilimonas*, part of *Chitinophagaceae*, are associated with degradation of complex polysaccharides like chitin and cellulose, indicating adaptation to slow but sustained substrate availability. Although little is known about *Palsa-955*, its consistent detection in undisturbed conditions suggests traits supporting persistence in stable environments. Among ruderals (R), *Sphingomonadaceae* are well known for degrading a wide range of xenobiotic compounds and for their metabolic flexibility, allowing fast adaptation to fluctuating substrates [47]. *Sumerlaeia* has been associated with nitrogen cycling in Antarctic soil ecosystems where microbiota responded rapidly to augmented nutrient regimes [48]. *OLB8* and *OLB17*, although poorly characterized [49], have been previously enriched in reactors under varied disturbance regimes [21], suggesting rapid growth and flexible substrate uptake as ruderal traits. Stress-tolerant (S) taxa include *Zoogloea*, which form protective extracellular matrices, are motile, and engage in aerobic denitrification, traits that could enhance survival in stressed environments [50]. Members of *Burkholderiaceae* are often capable of surviving in oligotrophic conditions and possess stress-response pathways such as DNA repair and oxidative stress resistance. *UBA2336* and *Runella*, though less studied, belong to lineages that are commonly found in activated sludge and have been linked to biofilm formation and resilience under harsh conditions [51]. These assignments support the use of MAG-level CSR classifications to interpret microbial succession and functional adaptation in bioreactors across disturbance gradients.

Trait-based CSR interpretation is gaining traction in both natural and engineered systems, with studies in soil showing microbial trade-offs that parallel Grime’s original framework [22, 29, 52, 53]. Our results extend this approach to activated sludge, demonstrating its relevance for bioreactors where microbial communities are managed for nutrient removal or resource recovery. By reducing microbial complexity into three ecologically meaningful axes, CSR offers a practical framework for integrating structure, function, and traits [2], while tools like MicroEcoTools streamline this process using reproducible metrics [37]. Other models, such as the Y-A-S framework (Yield, Acquisition, Stress tolerance), emphasize carbon-use efficiency and substrate uptake and have been applied in soils [54, 55]. Despite conceptual overlaps with CSR, such models may serve as complementary tools rather than alternatives [31].

Further work is needed to evaluate the broader applicability of CSR as a tool for microbial resource management, particularly in systems relevant to biotechnology such as anaerobic digestion [56], and at different operational and spatial scales, including systems with varying degrees of environmental control and bacterial immigration [2]. Controlled comparisons across systems will help assess the relative strengths and limitations of CSR and other frameworks. Our findings support CSR as a practical, interpretable approach for engineered microbial communities, with strong potential for broader integration alongside emerging ecological and predictive modelling tools.

## Conclusions

This study reinforces the utility of classical life-history theory as a unifying framework for interpreting microbial community dynamics across environmental contexts. By integrating taxonomic, functional, and genome-resolved trait data across a disturbance gradient, we show that strategy differentiation among competitors, ruderals, and stress-tolerants can be detected and meaningfully interpreted. The use of MAG-based community-aggregated traits improves trait resolution for assembled taxa, offering deeper ecological insights, although some relevant functions within the community may be missed due to assembly limitations. Machine learning approaches, such as DistLM, enhance this framework by identifying trait combinations that predictably explain community structure, contributing to the development of interpretable, data-driven models of microbial responses to environmental change. Open-source tools like MicroEcoTools enable scalable and reproducible classification of microbial communities, supporting wider application of trait-based approaches. The convergence of results from independent studies in activated sludge systems, each employing different disturbance types, further supports the generality of this framework. As microbial ecology increasingly aims to generate predictive, theory-grounded insights, such integrative approaches provide a bridge between classical ecological models and contemporary microbial genomics.

## Materials and Methods

### Experimental Design and Functional Assessments

This study builds upon the experiment previously described in <u>Santillan and Wuertz</u> [36], where detailed experimental procedures and baseline analyses are provided. Here, we focus on additional analyses and datasets not included in that publication. Briefly, we conducted a 42-day experiment using thirty microcosm-scale sequencing batch bioreactors (25 mL working volume), each inoculated with activated sludge from a full-scale municipal wastewater treatment plant in Singapore. Bioreactors were maintained at 30°C and operated in daily cycles. A synthetic medium was supplied following a fixed feeding regime with double organic loading applied at six disturbance frequencies: never (undisturbed), every 8, 6, 4, or 2 days (intermediate disturbance), and daily (press disturbance), each in five independent replicates (n = 5). These were labelled levels 0 to 5, corresponding to increasing disturbance frequency, with calculated frequencies of 0, 1/8, 1/6, 1/4, 1/2, and 1, respectively. System performance was monitored weekly by measuring soluble COD and TKN removal at the end of a cycle using spectrophotometric and ion chromatographic methods [57]. Settling capacity was assessed via SVI (mL/g), with 30 min of settling time. For correlation analyses, we used the additive inverse (−SVI) so that higher values reflect better settleability, as lower SVI values correspond to improved settling performance. Feed concentrations in mixed liquor after feeding were ∼306 mg COD/L and ∼46 mg TKN/L under regular conditions and ∼595 mg COD/L and ∼46 mg TKN/L under double loading. Biomass levels were controlled using a food-to-biomass (F:M) ratio strategy as described in <u>Santillan et al.</u> [13], targeting a total suspended solids (TSS) concentration of 1,500 mg/L. The corresponding solids residence times (SRTs) ranged from 30 to 15 days as disturbance frequency increased. These values remain within typical doubling time thresholds for activated sludge bacteria [58].

### Sludge Inoculum and Reactor Setup

The inoculum was sourced from a nitrifying-denitrifying activated sludge process with ∼200,000 m³/d flow and ∼80–90% nutrient removal efficiency. On the day of collection, influent COD and TKN concentrations averaged 221 mg/L and 45 mg/L, respectively. Activated sludge was transferred to the lab in sterile conditions, homogenized, and distributed into 30 sterile 50-mL tubes acting as sequencing batch reactors (SBRs). Each reactor was fed 12.5 mL of either regular or high-strength synthetic wastewater, depending on the disturbance schedule. A daily cycle involved 30 min of settling, decanting 12.5 mL of effluent, and replacing it with fresh medium. This setup yielded a hydraulic residence time (HRT) of 48 hours.

### Synthetic Wastewater Composition

The synthetic wastewater composition followed previous formulations <u>Santillan et al.</u> [13], including complex organics (yeast extract, peptones), simple carbon sources (acetate, dextrose), inorganic nitrogen sources (urea, ammonium salts), buffering agents, and trace elements. For double organic loading events, COD inputs were doubled primarily through the organic components, while inorganic nitrogen was reduced slightly to maintain TKN levels. Phosphate was added to target an N:P ratio of ∼6. Media were prepared in large batches, sterile-filtered (0.2 μm), and stored at 4°C.

### DNA Extraction and 16S rRNA Gene Amplicon Sequencing

Sludge samples were collected weekly from all reactors (n = 180), plus inoculum controls (n = 4), and stored at –80°C. Genomic DNA was extracted using the FastDNA Spin Kit for Soil (MP Biomedicals) with previously described protocol modifications to improve yield [21]. DNA was quantified using NanoDrop and Qubit instruments and further purified before sequencing.

The 16S rRNA gene V3–V4 regions were amplified using the 341f/785r primer set in a two-step PCR protocol [13, 59]. Dual-indexed libraries were generated, pooled, and sequenced using the Illumina MiSeq v3 platform with 300 bp paired-end reads. Amplicon sequence variants (ASVs) were inferred using the DADA2 pipeline [60], with quality filtering, error correction, merging, and chimera removal. Taxonomy was assigned with the SILVA database (v.132) [61]. Rarefaction was performed to 5,089 reads per sample to normalize sequencing depth for downstream diversity and community analyses.

### Metagenomics Bioinformatics

Metagenomic analyses followed a previously described pipeline [34], optimized for recovering and characterizing non-redundant, medium-to high-quality MAGs. Raw reads were trimmed using Trimmomatic [62] to remove adapters and low-quality bases (Q < 30), with FastQC [63] used for quality assessment. Reads from replicate and time-series samples were co-assembled using SPAdes [64] in metagenomics mode with multiple k-mer sizes. Coverage profiles were generated by mapping reads to contigs using BBMap [65], and genome binning was performed with MetaBAT2 [66]. Resulting bins were dereplicated using dRep [67], selecting species-level MAGs with ≥50% completeness and ≤10% contamination, assessed using CheckM [68].

Taxonomic classification was carried out with GTDB-Tk [69], and open reading frames (ORFs) were annotated with Prokka [70] and functionally classified via EggNOG-mapper. Ribosomal and transfer RNA genes were annotated using Barrnap [71] and tRNAscan-SE [72], respectively. MAG relative abundances were estimated with CoverM [73], using genome-based quantification with minimum coverage thresholds. This pipeline enabled robust recovery of MAGs and consistent functional annotation, supporting downstream trait-based ecological analyses.

### Community Structure, Genotypic Traits, and CSR Analysis

Community function, structure, and genotype data were analysed using the CSR framework as outlined in [21]. Community structure was assessed at the genus level using both metagenomics and 16S rRNA gene amplicon sequencing data, combining ordination methods and multivariate tests in PRIMER (v.7) [74]. Square-root transformed, normalized genus abundances were used to reduce the influence of dominant taxa. To evaluate links between community structure and function across disturbance levels, constrained ordination via canonical analysis of principal coordinates (CAP) was conducted, incorporating Pearson’s correlation vectors of normalized functional metrics. Community differences by disturbance level were tested using PERMANOVA on Bray-Curtis dissimilarity matrices [75], with factors treated as fixed. Multivariate dispersion homogeneity was assessed using PERMDISP [76], and all p-values were calculated with 9,999 permutations.

Relationships between genotype-level traits and community structural patterns were explored using distance-based linear modeling (DistLM) applied to MAG data, and distance-based redundancy analysis (dbRDA) was performed using COG trait-complexes as predictor variables [77]. Pearson’s correlation vectors (r > 0.20) were overlaid on the dbRDA plots. CSR life-history strategy assignments were derived from both taxonomic and genotypic data using the *CSR_assign()* function from the MicroEcoTools package [37]. Composition-aware network analysis of MAG genotypic profiles was performed using Gephi (v.0.9.2), based on SparCC correlation matrices generated with the *sparcc* function from the SpiecEasi R package (v.1.1.0) [78], following [11]. Weak correlations (r_adj_ < 0.2) were removed. Node clusters were defined by Louvain modularity class, with clusters colored by their association with undisturbed or disturbed conditions. Networks were visualized using the Fruchterman-Reingold layout; node size was scaled by degree, and edge thickness by correlation strength.

## Acknowledgements

This research was supported by the Singapore National Research Foundation and Ministry of Education under the Research Centre of Excellence Program. We thank JYJ Tan, JQ Teo, A Latiff, SS Thi, AFBM Batcha, CK Aw, ABA Aziz, and JHJ Lim for their assistance with laboratory work. We thank LCW Liew for supporting the collection of sludge inoculum.

## Author Contributions

ES and SW conceived the idea. ES designed and conducted the experiment, and performed data processing and analysis. SAN carried out the metagenomic bioinformatics. SW secured funding for the study. ES drafted the manuscript, and all authors contributed to its revision and editing.

## Data availability

DNA sequencing data are available in the NCBI BioProject database under accession number PRJNA723443. A set of dereplicated MAGs at the genus level is available via GenBank under BioProject accession number PRJNA108977245. Additional files, including strain-level dereplicated MAGs (FASTA), functional annotations, relative abundances across samples, and phylogenetic tree files, are available through Zenodo (DOI: 10.5281/zenodo.8405311), as described in [34]. Summary statistics, taxonomic classifications, quality metrics, and MicroEcoTools *CSR_assign* classifications for the MAGs are available online in Supplementary Table S3.

