## Supplementary Information for "Predicting microbial community responses to disturbance using genome-resolved trait-based life-history strategies"

*Ezequiel Santillan^1^, Soheil Asgari Neshat^1^, and Stefan Wuertz^1,2*^*

^1^Singapore Centre for Environmental Life Sciences Engineering, Nanyang Technological University, Singapore, 637551, Singapore.

^2^School of Civil and Environmental Engineering, Nanyang Technological University, Singapore, 639798, Singapore.

**Table S1.** Community-level process performance data on day 42.

| **Community function**^†^ | **Disturbance frequency levels^*^** | | | | | | **Welch’s ANOVA**  **P_BH_**^§^ |
| --- | --- | --- | --- | --- | --- | --- | --- |
|  | **0** | **1** | **2** | **3** | **4** | **5** |  |
| COD rem^¶^ (%) | 98.2 (1.3) | 98.1 (0.7) | 99.1 (0.7) | 98.7 (0.4) | 98.7 (0.2) | 98.6 (0.3) | 0.41 |
| TKN rem^¶^ (%) | 95.5 (2.5) | 97 (3.2) | 97.3 (0.5) | 97.8 (0.5) | 96.1 (1.7) | 54.6 (9.2) | **<0.001** |
| TKN | 2.06 (1.14) | 1.36 (1.49) | 1.22 (0.22) | 1.04 (0.23) | 1.8 (0.77) | 21.0 (4.25) | **<0.001** |
| NO_2_⁻-N [mg/L] | 3.66 (5.54) | 3.9 (2.08) | 0.82 (0.27) | 2.18 (0.61) | 2.78 (0.16) | 0 (0) | **<0.001** |
| NO_3_⁻-N [mg/L] | 54.4 (6.09) | 23.1 (8.05) | 52.3 (2.65) | 25.7 (3.65) | 15.2 (1.09) | 0 (0) | **<0.001** |
| PO_4_^3^⁻-P [g/L] | 8.28 (0.37) | 7.3 (0.54) | 8.96 (0.40) | 7.82 (0.45) | 5.82 (0.46) | 10.1 (0.26) | **<0.001** |
| SO_4_^2^⁻-S [g/L] | 26.4 (0.30) | 26.7 (1.01) | 26.4 (0.34) | 27.8 (1.70) | 28.0 (1.83) | 28.0 (2.17) | 0.26 |
| TA | 36.4 (13.2) | 177 (28.5) | 54 (9.5) | 173 (15.2) | 224 (3.00) | 468 (16.9) | **<0.001** |
| SVI | 56.4 (7.2) | 45.4 (6.9) | 39.8 (4.7) | 37.6 (4.0) | 39.2 (4.5) | 43.4 (2.5) | **0.017** |
| VSS:TSS (%) | 88.6 (2.9) | 94.4 (4.2) | 93.6 (1.7) | 91.6 (2.3) | 92.6 (0.9) | 93.8 (3.0) | 0.16 |

^†^ COD, chemical oxygen demand; TKN, total Kjeldahl nitrogen; TA, total alkalinity; SVI, sludge volume index; VSS:TSS, volatile to total suspended solids ratio.

^*^ Average values (n = 5), with standard deviation in parentheses.

^¶^ Removal of the indicated compound (percentage based on feed input).

^§^ Welch’s ANOVA test, Benjamini-Hochberg corrected P-value (significant values in bold).

**Table S2**. Quality assessment of 133 medium- and high-quality metagenome-assembled genomes (MAGs) recovered from day-42 bioreactor metagenomes.

| **Completeness^†^** | **Range** | **Count** |  | **Contamination**^*^ | **Range** | **Count** |
| --- | --- | --- | --- | --- | --- | --- |
| moderate | 50 - 69.9 % | 27 |  | medium | 5 - 9.9 % | 28 |
| near | 70 - 89.9 % | 49 |  | low | 0 - 4.9 % | 91 |
| substantial | 90 - 99.9 % | 55 |  | none | 0 % | 14 |
| perfect | 100 % | 2 |  |  |  |  |

**^†^**Completeness categories based on the Minimum Information about Metagenome-Assembled Genomes (MIMAG) framework [38].

^*^Contamination categories and their corresponding counts.
